# Gain and loss of function changes in *CACNA1C* affect neuronal networks through divergent pathways

**DOI:** 10.64898/2025.12.12.693190

**Authors:** Gemma Wilkinson, Jamie Wood, Nuppu Roivainen, Jack Underwood, Jeremy Hall, Adrian J Harwood

## Abstract

**Background:** *CACNA1C* encodes the pore-forming subunit of the L-type calcium channel Cav1.2. Common variants in CACNA1C are associated with psychiatric disorders, while rare single-nucleotide variants cause CACNA1C-related disorder (CRD), a multi-system disorder with symptoms that include autism spectrum disorder (ASD), intellectual disability and seizures. However, the cellular mechanisms linking CACNA1C dysfunction to neurodevelopmental phenotypes remain poorly understood.

**Methods:** We generated isogenic *CACNA1C* loss-of-function (LoF) induced pluripotent stem cell (iPSC) lines and reprogrammed a line from an individual carrying a novel gain-of-function (GoF) variant in *CACNA1C*. Neuronal activity was assessed using multi-electrode arrays (MEA), pharmacological manipulation, and gene expression analysis. Early developmental phenotypes were examined using qRT-PCR, immunocytochemistry, and RNA sequencing.

**Results:** LoF and GoF patient-derived neurons displayed opposing alterations in network dynamics. Pharmacological and molecular assays indicated that these network differences were associated with dysregulated GABAergic signalling. Early developmental analysis revealed that loss of *CACNA1C* altered rosette morphology, CREB phosphorylation, and transcriptional programs related to axonogenesis and synaptic signalling, indicating effects on neuronal differentiation. The patient line exhibited opposing phenotypes, reflecting variant-specific effects.

**Conclusions:** These findings demonstrate that Cav1.2 regulates excitatory–inhibitory balance, network organization, and aspects of neurodevelopment. Divergent effects of LoF and GoF variants highlight how altered Cav1.2 signalling contributes to variable neurodevelopmental phenotypes, including ASD and epilepsy, and establish a framework for defining CACNA1C variant effects in human neurons.

## Introduction

The L-type calcium channel (LTCC) Cav1.2, encoded by *CACNA1C*, has been strongly associated with psychiatric disorders including bipolar disorder, schizophrenia and autism spectrum disorder via genome wide association studies (GWAS)^1–4^. In addition to this, patients with variants in *CACNA1C* present with a range of neurodevelopmental symptoms. The most well-studied is the gain-of-function variant G406R, which causes Timothy Syndrome (TS), a multi-system disorder characterised by syndactyly, congenital heart disease, long QT syndrome, developmental delay, autism spectrum disorder (ASD) and seizures^5^. Patients without the core features of Timothy Syndrome, often with novel variants in *CACNA1C* are now collectively termed as *CACNA1C*-related disorder (CRD) ^6^. These patients exhibit variable phenotypes: some patients show primarily cardiac symptoms, while others present with more prominent neurodevelopmental or neurological features^7,8^. To date, only the canonical G406R TS variant has been studied in neuronal models, where it has demonstrated substantial effects on neurogenesis, neuroplasticity and axonal maturation^9–13^. However, little is known about how different *CACNA1C* variants influence neuronal development and function.

Cav1.2 plays an important role in shaping neuronal activity. It regulates synaptic plasticity via long term potentiation (LTP) with deletion in rodents leading to a reduction in LTP and selective memory deficits^14–16^. At the single-cell level Cav1.2 regulates action potential dynamics, and prolonged opening of Cav1.2 channels in TS neurons leads to extended depolarisation and wider action potentials^9^. At the network level, LTCCs regulate oscillatory activity. Synchronised calcium transients in cultured neurons are reduced LTCC blockade and LTCC activation is required for theta oscillations^17–19^. CACNA1C, specifically has been shown to be important in phase coupling of theta and gamma oscillations^16^. Finally, in iPSC-derived neurons blockade of LTCCs alters the pattern of synchronised burst activity by reducing the interval between bursts^20^. The formation of functional neuronal networks is a key aspect of brain development and synchronised activity has is an important intrinsic property of neurons^21–25^. These network-level effects are likely relevant to neuropsychiatric symptoms associated with *CACNA1C*, including ASD and seizures.

Despite the clear effects of LTCCs on network activity, many of the findings come from pharmacological studies, which cannot distinguish between Cav1.2 and the Cav1.3 channels, making it difficult to attribute network-level effects to a specific LTCC subtype. As a result, the specific contribution of Cav1.2 to the formation and synchrony of neuronal networks has not been directly investigated in human cellular models. Moreover, the mechanisms linking Cav1.2 function to network excitability remain poorly understood. LTCCs also regulate cortical development and altered progenitor differentiation or excitatory–inhibitory balance may affect network formation^9,26,27^. Understanding how Cav1.2 shapes neuronal network activity could reveal critical insights into the neurophysiological basis of *CACNA1C*-related disorders.

To better understand the role of *CACNA1C*, we investigated iPSC-derived neurons from cell lines with a *CACNA1C* loss of function (LoF) variant and from a patient with a novel GoF *CACNA1C* variant. Using MEA recordings and pharmacological manipulation of LTCCs and GABAergic signalling, we demonstrate that *CACNA1C* critically regulates synchronised neuronal network activity. These findings highlight a potential mechanism by which *CACNA1C* variants contribute to neurodevelopmental and neurological symptoms and suggest avenues for therapeutic intervention.

## Methods and Materials

Full details of experimental procedures can be found in Supplemental Methods

### Cell Lines

CACNA1C knockout (LoF) lines were generated via CRISPR-Cas9 targeting exon 2 in an inducible Cas9 iPSC line. A patient with a *CACNA1C* p.Ala1521Pro variant was recruited via the IMAGINE-ID study and 2 iPSC clones were generated from a peripheral whole-blood sample. Phenotypic features of the individual included autism, developmental delay and intellectual disability, borderline cardiac QT interval and muscular hypotonia^7^. 3 control lines were used in the study; an IBJ4 human iPSC line derived from the BJ fibroblast cell line, HPSI1013i-wuye_2 and an inducible Cas9 line, which was derived from the IBJ4 line.

### Neuronal differentiation

Differentiation into cortical glutamatergic neurons followed a modified protocol by Chambers et al. using dual SMAD inhibition (SB431542, LDN193189) in N2B27 media^28^. At day 50 of differentiation, neurons were plated onto multi-electrode arrays (MEAs) for electrophysiological assays and maintained in BrainPyhs medium supplemented with B27+RA, BDNF and ascorbic acid.

### MEA recordings and pharmacology

Neuronal activity was recorded using an Axion Maestro on 24-well MEA plate. Spike detection and synchrony analyses were performed using AxiS software and custom R scripts. Pharmacological manipulations included bicuculline, diltiazem, and diazepam, applied acutely. Wells with less than 80% electrode coverage were excluded from analysis.

### RNA sequencing

RNA sequencing was performed on day 30 neurons (n = 3 per genotype). Libraries were prepared and sequenced by Active Motif. Reads were aligned using STAR, and differential expression was analyzed in R (DESeq2, v1.30.0) with Benjamini–Hochberg correction (adjusted P < 0.01, |log₂FC| ≥ 0.5). Functional enrichment was assessed using fgsea, with clustering of Gene Ontology terms via GOSemSim. Tissue and disease enrichment analyses used TissueEnrich and DisGeNET, respectively.

### Protein and immunocytochemical analysis

Western blotting and immunocytochemistry were performed using standard protocols. Protein expression of Cav1.2 and downstream signalling targets (ERK1/2, CREB) was analysed by western blot using infrared detection. Immunostaining for neuronal and progenitor markers (Pax6, Ki67, NeuN) was performed on fixed cells, imaged using a Leica DMI6000b microscope, and quantified with CellProfiler.

### Quantitative PCR

RNA was extracted using the RNeasy kit, reverse transcribed with the High-Capacity cDNA kit, and analysed by qPCR using the Pfaffl method, normalized to GAPDH and β-actin.

### Statistical Analysis

Statistical analyses were performed in GraphPad Prism 8. Normality was assessed using the Anderson–Darling test, and one- or two-way ANOVAs were applied as appropriate. Data are presented as mean ± SEM, with significance thresholds of P < 0.05.

## Results

### LTCC blockade reduces excitability

Previous studies have shown that acute pharmacological blockade of LTCCs leads to changes in neuronal network activity in both rodent and human cell models^17,18,20^. Our initial experiment aimed to confirm these findings in our human iPSC-derived cortical neurons. As previously seen, control neurons with wild-type CACNA1C genes displayed synchronised network activity with repeated periods of bursting behaviour (Fig. 1a). When the LTCC antagonist diltiazem was applied to neurons we found a significant reduction in neuronal excitability, demonstrated by a decrease in the well spike rate (Fig. 1b) and the number of active electrodes (Fig. 1c). Consistent with the previous studies, we also showed that diltiazem led to a significant reduction in the interval between synchronised bursts (SBs) (Fig. 1d). There was no effect on the percentage of spikes occurring within SBs (Fig. 1e). This measure shows there is no change in the level of synchronicity of cultures following diltiazem, even though the frequency of bursts is altered.

**Figure 1:**
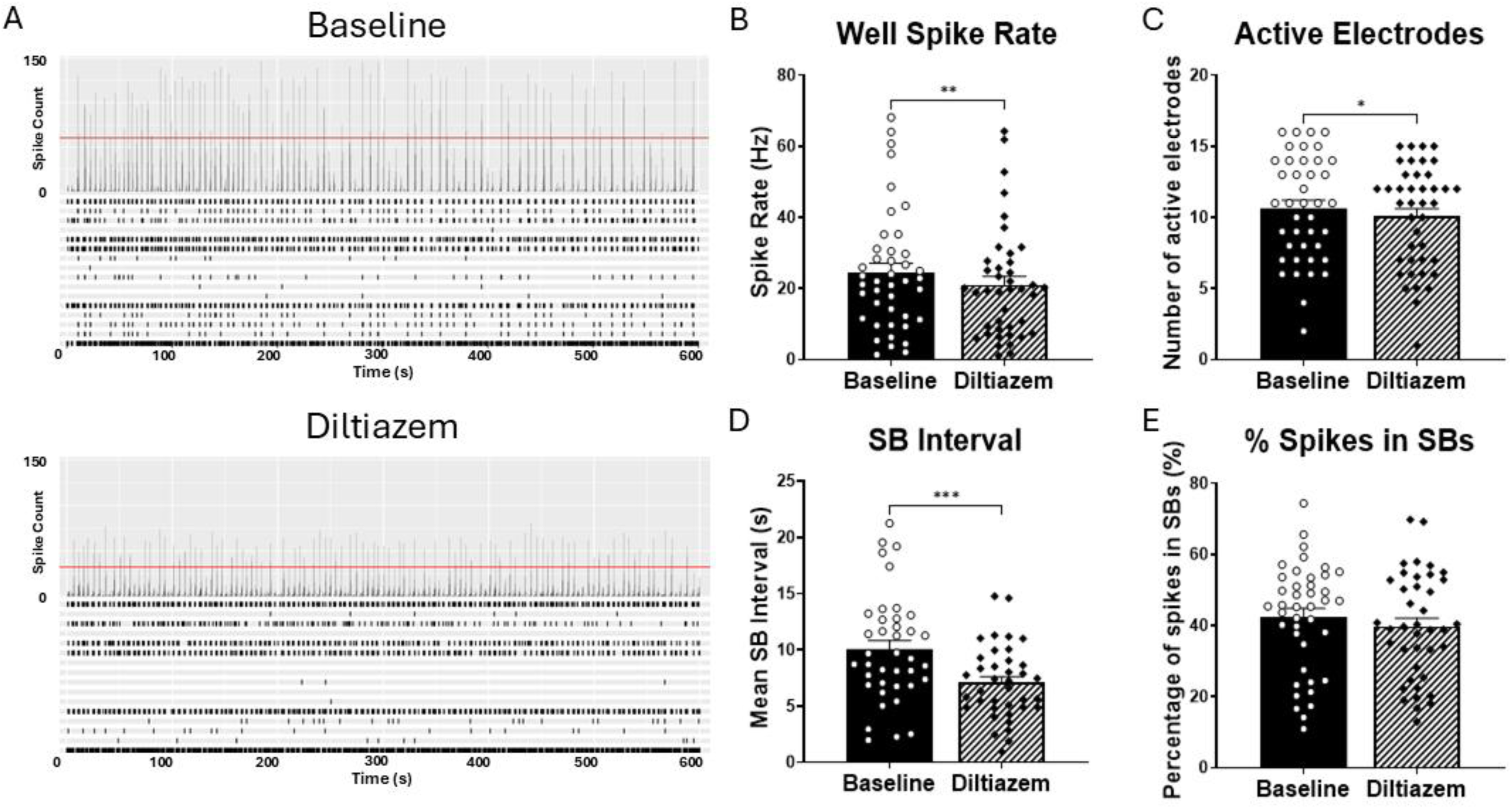
Effect of LTCC blockade on neuronal network activity. A: Representative raster plots with array-wide spike detection in control neurons at baseline and following diltiazem treatment. For both conditions, the upper panel shows the number of spikes per 200ms bin across a 600 second recording. Vertical scale = 200 spikes per bin. The lower panel shows a raster plot, each row corresponds to an electrode and spikes are plotted as black lines across the 600 second recording. B-C: Basal excitability is shown by well spike rate and number of active electrodes. D-E: Measures of synchronised activity are mean interval between synchronised bursts (SBs) and percentage of spikes occurring within SBs across the 600 second recording. All plots show mean ± SEM, individual wells are overlayed as points. *p < 0.05, **p < 0.01, *** p < 0.001, **** p < 0.0001 following paired T-test. Data from 3 independent differentiations across 3 control cell lines.

### CACNA1C variants oppositely affect network activity

To identify the role of *CACNA1C* specifically on neuronal activity we generated iPSC-derived neurons from a patient carrying a novel variant in *CACNA1C* alongside two LoF lines. The LoF lines were generated using CRISPR-Cas9, which introduced deletions in exon 2 of *CACNA1C* leading to a frameshift and introduction of a stop codon in exon 3 (Fig. S1b-e). Amplification of RNA of exons 1-4 of *CACNA1C* transcript demonstrated that these were homozygous mutants as no wild-type message was found (Fig. S1c). However, both lines only showed a 75% reduction of detectable Ca1.2 protein (FigS1f). This residual protein may arise due to translation reinitiation, possibly from the next methionine (p.Met265) in exon 6 of *CACNA1C* (NM_000719.7) and therefore is unlikely to be functional. The iPSCs derived from our patient line contain a novel single nucleotide variant in *CACNA1C*; p.Ala1521Pro in exon 38 (Fig. S1a). This variant has not previously been tested but is predicted to be deleterious by two independent pathogenicity predictor algorithms. PolyPhen-2 predicted the variant to be probably damaging with a score of 1.0 out of 1.0. Provean predicted itto be deleterious with a median score of −4.807, scores equal to −2.5 or below are considered deleterious^29,30^. Western blot showed that this variant had no effect on protein expression of Cav1.2 in neurons (Fig. S1g). This suggests any phenotypes seen in this cell line were due to changes in channel function, rather than altered expression.

Recordings of spontaneous activity using multi-electrode arrays (MEA) at day 100 showed that LoF and patient neurons formed synchronised network behaviour (Fig. 2a). However, the two exhibited distinct and opposing effects, suggesting a gain of function (GoF) effect on Cav1.2 activity in the patient line. Patient (GoF) neurons exhibited a significant reduction in excitability, evidenced by decreased well spike rate, fewer active electrodes, and reduced spike amplitude (Fig. 2b-c, Table S2). In contrast, LoF lines showed no significant changes in spike rate. However, they did demonstrate a significant increase in the number of active electrodes, suggesting an elevated excitability. LoF and GoF lines also exhibited opposing changes in synchronized burst interval and burst frequency (Fig. 2d). Specifically, LoF lines displayed more frequent synchronized bursts with shorter intervals between them, whereas GoF lines had fewer bursts with longer intervals. The GoF lines also showed a reduction in spikes per synchronized burst, likely reflecting the overall decrease in spike rate (Table S2). LoF lines showed no significant change in spikes per burst but had an increased percentage of spikes occurring within synchronized bursts, indicating enhanced network synchrony (Fig. 2e).

**Figure 2:**
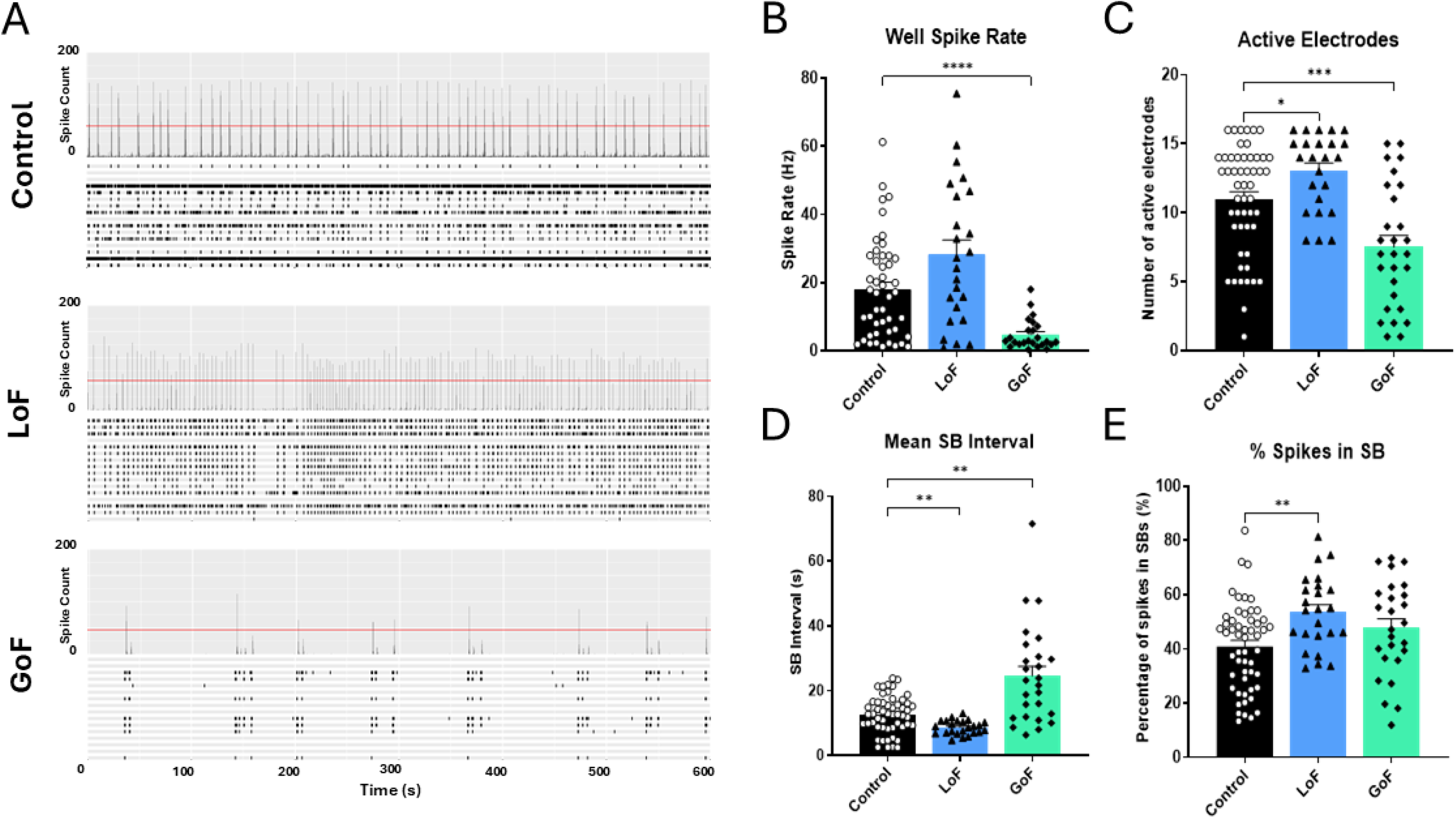
Changes to CACNA1C alter neuronal network activity. A: Representative raster plots with array-wide spike detection in control, LoF and GoF neurons. For each group, the upper panel shows the number of spikes per 200ms bin across a 600 second recording. Vertical scale = 200 spikes per bin. The lower panel shows a raster plot, each row corresponds to an electrode and spikes are plotted as black lines across the 600 second recording. B-C: Basal excitability is shown by well spike rate and number of active electrodes. D-E: Measures of synchronised activity are mean interval between synchronised bursts (SBs) and percentage of spikes occurring within SBs across the 600 second recording. All plots show mean ± SEM, individual wells are overlayed as points. *p < 0.05, **p < 0.01, *** p < 0.001, **** p < 0.0001 following One Way ANOVA. Data from at least 3 independent differentiations across 2-3 cell lines per group.

### GABAergic signalling shapes network phenotypes

It is well established that GABA plays an important role in controlling network activity, including the interval between synchronised bursts^31,32^. Therefore, we decided to target GABAergic signalling to see if we could rescue the changes we saw in the *CACNA1C* mutant lines. In the LoF neurons, we applied diazepam, a GABAA positive allosteric modulator (Fig. 3a). Following baseline recordings, 50 nM diazepam was applied for 1 hour. At baseline, LoF neurons had an elevated spike rate compared to controls and this was also present post-diazepam treatment (Fig. 3b). Diazepam led to a significant reduction in spike rate in the control neurons but had no effect in the LoF. Similar results were observed for the number of active electrodes, which was significantly higher in LoF neurons at both baseline and after diazepam treatment (Fig. 3c). Again, diazepam significantly reduced this measure only in controls. Diazepam treatment had no significant effect on synchronised burst interval in either control or LoF neurons (Fig. 3d). However, there was a significant reduction in SB interval in LoF neurons compared to controls, replicating the findings of the previous experiment. Post-hoc tests showed that this approached significance at baseline and was significant following diazepam treatment. The percentage of spikes occurring in synchronised bursts was also elevated in LoF neurons at both timepoints, and although there was a main effect of treatment, post-hoc tests indicated that only control lines exhibited a significant reduction in synchronised bursting (Fig. 3e). Overall, diazepam reduced network excitability and synchrony, but these effects were limited to the control lines, suggesting resistance to GABAergic modulation in LoF cultures.

**Figure 3:**
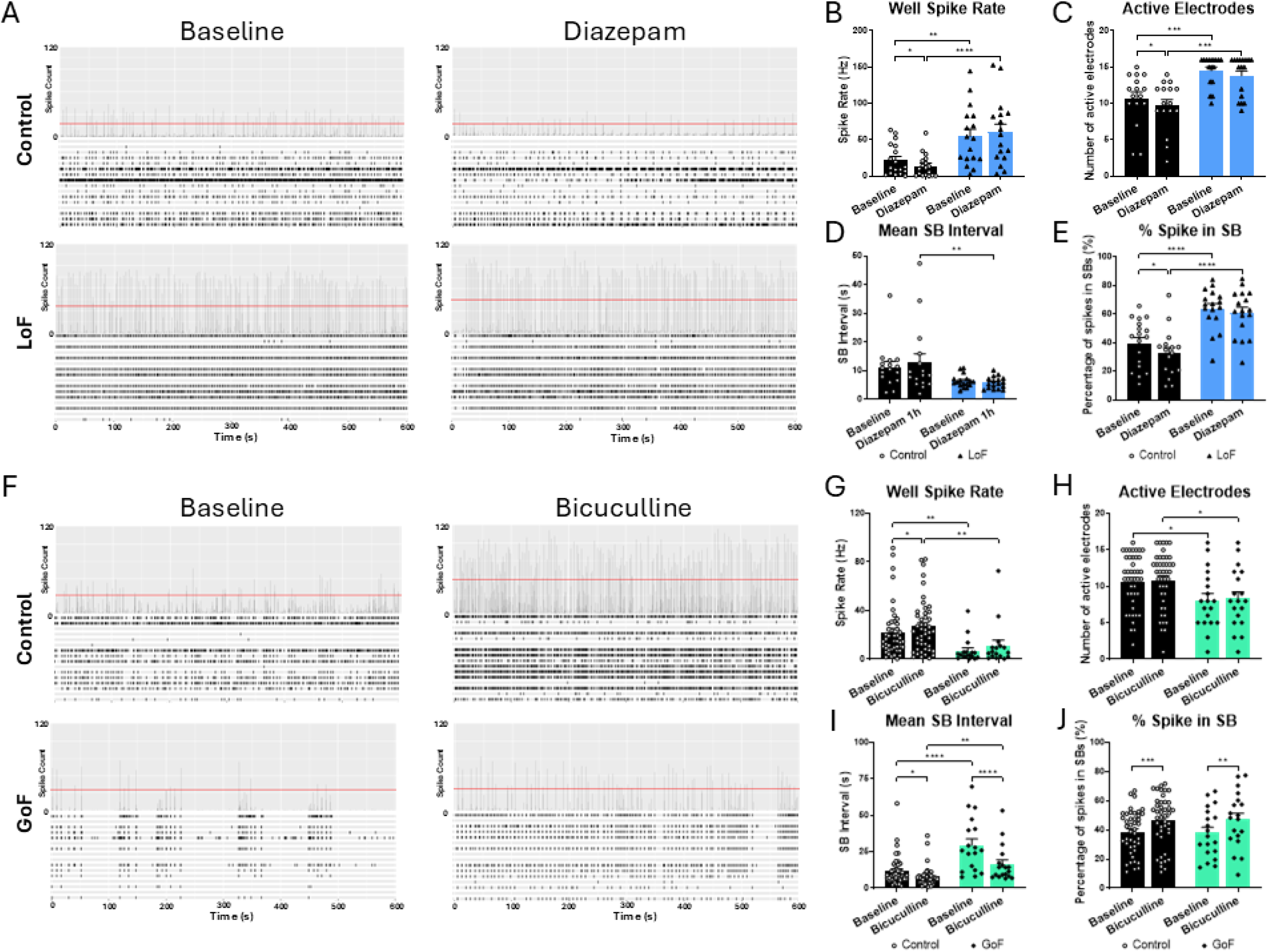
Pharmacological targeting of GABAA receptors in CACNA1C LoF and GoF lines. A: Representative raster plots with array-wide spike detection in control and CACNA1C LoF lines at baseline and following Diazepam treatment. For each condition, the upper panel shows the number of spikes per 200ms bin across a 600 second recording. Vertical scale = 200 spikes per bin. The lower panel shows a raster plot; each row corresponds to an electrode and spikes are plotted as black lines across the 600 second recording. B-E: Quantification of basal excitability and synchronised activity is shown by well spike rate, number of active electrodes, mean interval between SBs, and percentage of spikes occurring within SBs at baseline and following diazepam treatment in control and CACNA1C LoF lines. F: Representative raster plots with array-wide spike detection in control and CACNA1C GoF lines at baseline and following Bicuculline treatment. G-J: Quantification of basal excitability and synchronised activity is shown by well spike rate, number of active electrodes, mean interval between SBs, and percentage of spikes occurring within SBs at baseline and following bicuculline treatment in control and CACNA1C GoF lines. All plots show mean ± SEM, individual wells are overlayed as points. *p < 0.05, **p < 0.01, *** p < 0.001, **** p < 0.0001 following Two-Way ANOVA. Data from at least 3 independent differentiations across 2-3 cell lines per group.

In GoF neurons, we applied bicuculline, a GABAA receptor antagonist. 10 μM bicuculline was applied for 10 minutes following baseline recording (Fig. 3f). Consistent with previous results, spike rate was significantly reduced in GoF neurons compared to controls at both baseline and post-treatment (Fig. 3g). Bicuculline application led to a significant increase in spike rate, with a stronger effect in controls than GoF. The number of active electrodes was also significantly lower in GoF neurons, consistent with prior findings (Fig. 3h). However, bicuculline had no significant effect on this measure. For the mean SB interval, there were significant main effects of group and treatment, as well as a significant interaction (Fig. 3i). Post-hoc tests showed prolonged SB intervals in GoF neurons at both baseline and after treatment. Bicuculline significantly shortened the SB interval in both controls and GoF neurons. While there were no group differences in the percentage of spikes within synchronised bursts at baseline, bicuculline treatment led to a significant increase in synchrony in both control and patient neurons (Fig. 3j). Application of bicuculline partially reversed the hypoactive phenotype in GoF neurons, showing a strong effect on synchronised activity. However, its impact on basal excitability was limited in GoF neurons compared to controls.

### Excitatory-inhibitory imbalance underlies altered activity

Based on the effects of GABAergic drugs on the neuronal activity of the lines, we next investigated expression of genes important in GABAergic signalling. GABRA1 and GABRB2 encode the alpha1 and beta2 subunits of GABAA receptors and form the most abundant GABAA isoform. *CACNA1C* LoF neurons showed a significant reduction in GABRA1, although there was no significant change GoF neurons (Fig. 4a). Both GoF and LoF neurons showed a significant increase in GABRB2, with a more pronounced increase in the GoF (Fig. 4b). GAD65 and GAD67 encode the enzymes responsible for producing GABA. LoF neurons showed a significant reduction in both GAD65 and GAD67, whereas GoF neurons showed no significant changes (Fig. 4c-d).

**Figure 4:**
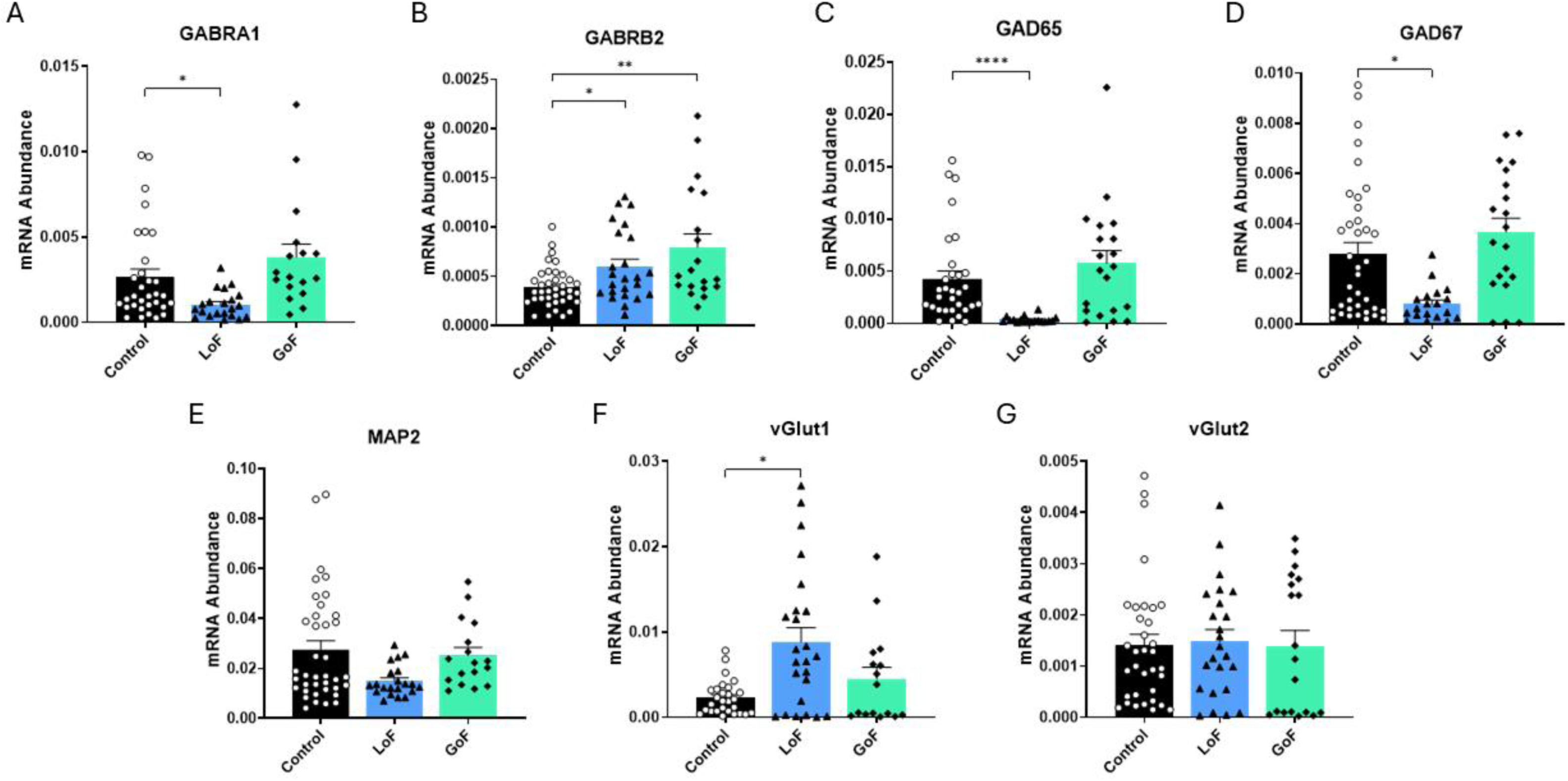
Expression analysis of genes encoding neuronal signalling components in control, CACNA1C LoF and CACNA1C GoF neurons at day 120. A-D: Analysis of a panel of inhibitory GABA signalling genes (GABRA1, GABRB2, GAD65, GAD67). E: Analysis of neuronal marker MAP2. F-G: Analysis of glutamate transporters as markers of excitatory neurons (vGlut1, vGlut2). The values are presented as relative mRNA abundance compared to expression of housekeeping genes GAPDH and ϐ.-Actin. All plots show mean ± SEM, individual wells are overlayed as points. *p < 0.05, **p < 0.01, *** p < 0.001, **** p < 0.0001 following One Way ANOVA. Data from at least 3 independent differentiations across 2-3 cell lines per group.

We also assessed MAP2, a general neuronal marker, LoF neurons trended toward lower MAP2 expression; and although the Kruskal-Wallis test indicated a significant group difference, post-hoc comparisons with controls were not significant (Fig. 4e). Next, we evaluated glutamate transporters vGlut1 and vGlut2 (Fig. 4f-g). LoF neurons showed increased vGlut1 expression, while vGlut2 levels remained unchanged across groups, consistent with its lower abundance in mature neurons. Altogether, these transcriptional changes suggest an altered excitatory-inhibitory balance, particularly in the LoF lines, that may underlie the observed neuronal activity changes and resistance to diazepam treatment.

### Transcriptional changes suggest developmental contribution

To explore the origins of the network phenotype we examined the early stages of neurogenesis with RNA-seq of day 30 samples, just as neurons are forming, in control and LoF lines. Differential expression analysis identified 142 upregulated and 128 downregulated genes compared to controls (Fig. 5a, Table S3). Upregulated genes were enriched for Gene Ontology (GO) biological processes related to axon development, behaviour, trans-synaptic signalling, synapse assembly, and regulation of neuron differentiation (Fig. 5b-c, Table S4). In contrast, downregulated genes were enriched for terms associated with forebrain neuron generation, cell junction organization, cell-cell adhesion, and regulation of transmembrane receptor protein serine/threonine kinase signalling.

**Figure 5:**
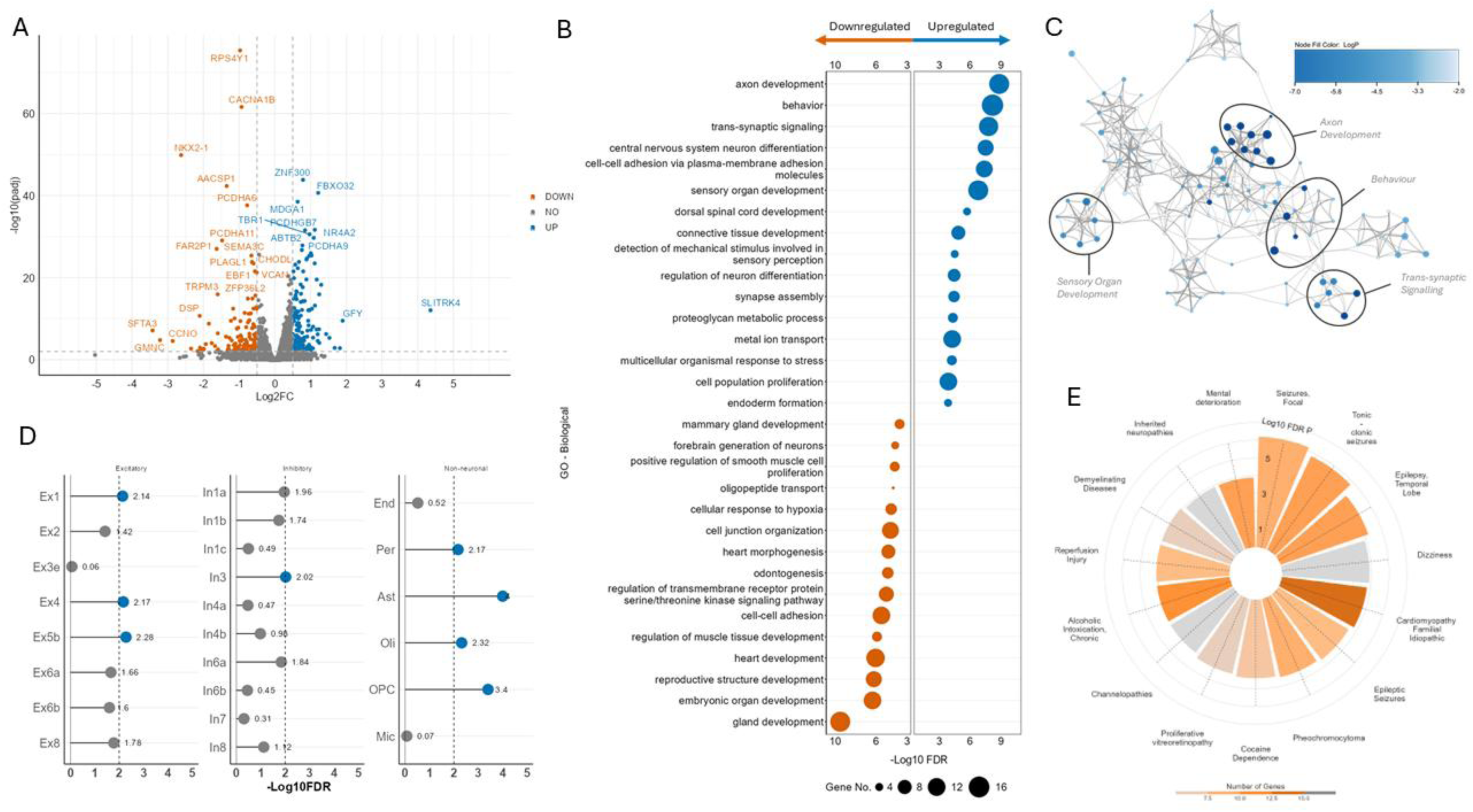
Differential gene expression of CACNA1C LoF cell line at day 30 compared to isogenic control cell line: A: RNAseq volcano plots of DEGs in a CACNA1C LoF mutant compared to wild type, determined by the DESeq2 method with a 0.5-fold change (FC) threshold and a false discovery rate (FDR)–adjusted P < 0.01 B: Ranked enrichment analysis of CACNA1C LoF gene expression signatures using the GOBP database by the fgsea multilevel enrichment test, genes ranked by minimum significant difference signed msd (fgsea). C: Network of enriched GOBP terms, clustered by shared ID and coloured by p-value, for upregulated DEGs. Enriched GOBP terms were clustered based on semantic similarity using GOSemSim (R package, v2.24.0), and clusters were manually summarized. Node size indicates the number of genes. D: Brain cell type enrichment analysis using gene expression per tissue based on brain tissue single cell data. Significant cell types P>0.05, are in blue. E Disease-gene enrichment analysis performed for all DEGs and disease associations from DisGeNET, with significance determined as P < 0.001. Bar plot colour indicates the number of enriched genes within a given disease gene-set.

Cell type enrichment analysis of these data revealed increased transcription of genes associated with the EX1, 4 and 5b subset of excitatory neurons, but no substantial enrichment for inhibitory neurons (Fig. 5d). Consistent with this, the interneuron transcription factors NKX2.1 and LHX6 were significantly downregulated, alongside LGI2, a gene required for inhibitory synapse formation (Table S3). These findings suggest an imbalance in the formation and function of excitatory and inhibitory neuron populations, consistent with the functional changes observed in MEA experiments. We also observed enrichment of astrocyte and oligodendrocyte gene signatures, suggesting reduced efficiency of neuronal differentiation, accompanied by a relative increase in glial cell generation within the cultures. Finally, disease association analysis highlighted strong links between the *CACNA1C* LoF gene signature and epilepsy-related phenotypes. Of the 14 diseases linked to *CACNA1C* loss, four were related to epilepsy and seizures, with focal seizures showing the strongest association (Fig. 5e). This aligns with our MEA findings of altered burst dynamics and network excitability, and increased seizure risk in patients with *CACNA1C* variants.

### Early NPC signalling is altered by *CACNA1C* variants

To determine whether the neuronal and glial cell-specific genes may arise due to changes in earlier development, prior to the emergence of distinct cell types, we examined signalling in neural progenitor cells (NPCs, day 20) and immature neurons (day 30). CREB phosphorylation, a downstream pathway of LTCCs and a regulator of NPC-to-neuron differentiation, showed opposing effects in LoF and GoF lines: pCREB was elevated in GoF NPCs at day 20 but reduced in LoF neurons at day 30 (Fig. 1a–b). These time-dependent, divergent changes mirror the opposite effects of LoF and GoF variants on network excitability.

We then assessed NPC maintenance and rosette organisation. The was no difference in expression of Pax6 expression in LoF and GoF lines at day 20 and day 30 (Fig. S2a-d). Ki67 expression was also unchanged at either timepoint (Fig. 6c-d, g-h), suggesting no gross differences in progenitor specification or proliferation. In contrast, N-Cadherin staining was altered in both LoF and GoF lines at day 20. While there were no changes in the number of rosettes present in cultures, the mean area of rosettes was altered with LoF lines showing smaller rosettes and GoF lines showing larger ones (Fig. 6e-f). Although this difference was lost by day 30 due to overall rosette decline, GoF lines retained more rosettes at this stage (Fig. 6i–j), consistent with prolonged NPC maintenance. As a large amount of neurogenesis should occur between day 20 and 30, we also investigated the expression of NeuN at day 30. LoF lines showed a significant decrease in the percentage of NeuN positive cells, suggesting a reduction in number of neurons, with no significant changes in GoF lines (Fig. S2e). The absence of changes in Pax6 or Ki67 at this timepoint, along with the enrichment of astrocyte and oligodendrocyte genes in the RNA-seq suggests the emergence of alternative, non-neuronal cell types in LoF cultures. These findings point to distinct effects of *CACNA1C* loss and variants on early neurodevelopment, which may contribute to later differences in network activity.

**Figure 6:**
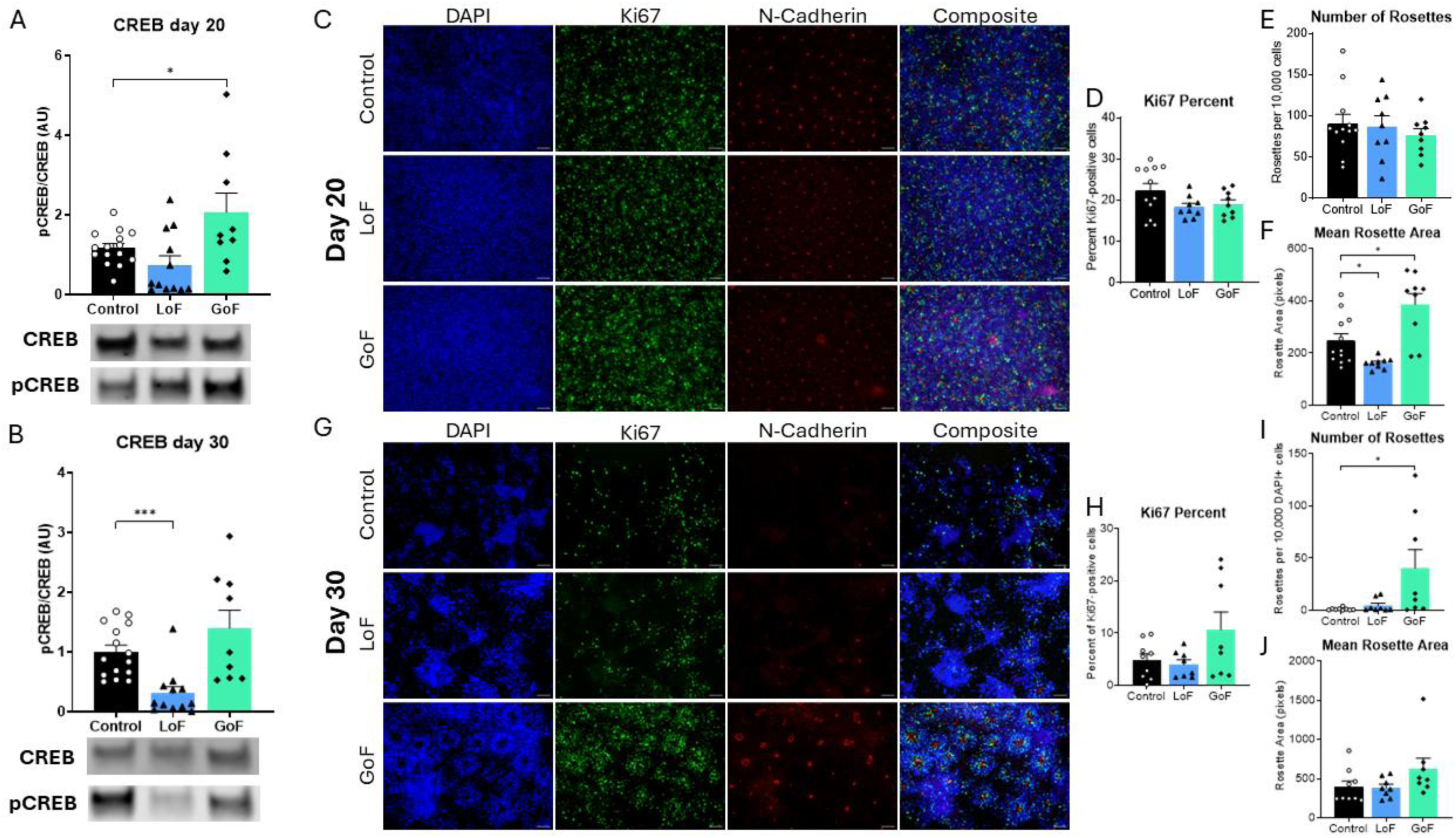
Altered neuronal differentiation of CACNA1C LoF and GoF lines. A-B: Phosphorylation of CREB at day 20 and day 30 of differentiation in control, CACNA1C LoF and GoF lines. CREB phosphorylation is measured as pCREB divided by total CREB expression. C: Examples images of Ki67 and N-Cadherin staining at day 20 of neuronal differentiation in control, CACNA1C LoF and GoF lines. D: Quantification of the percentage of DAPI+ nuclei that colocalized with Ki67 at day 20. E: Quantification of number of N-Cadherin+ structures normalized to 10,000 DAPI+ nuclei at day 20. F: Mean area of N-Cadherin structures at day 20. G: Examples images of Ki67 and N-Cadherin staining at day 30 of neuronal differentiation in control, CACNA1C LoF and GoF lines. H: Quantification of the percentage of DAPI+ nuclei that colocalized with Ki67 at day 30. I: Quantification of number of N-Cadherin+ structures normalized to 10,000 DAPI+ nuclei at day 30. J: Mean area of N-Cadherin structures at day 30. All plots show mean ± SEM, individual replicates are overlayed as points. *p < 0.05, **p < 0.01, *** p < 0.001 following One Way ANOVA. Data from at least 3 independent differentiations across 2 cell lines per group.

## Discussion

Our results show that *CACNA1C* is a key regulator of neuronal network activity in human iPSC-derived cortical neurons, with opposing effects depending on the type of variant. These network phenotypes were closely tied to altered GABAergic signalling, as pharmacological modulation partially rescued or revealed resistance in the mutant lines. Early changes in CREB signalling, NPC maintenance, and neuronal differentiation suggest that developmental mechanisms contribute to these later network differences, linking transcriptional and morphological alterations to functional outcomes. These findings highlight how *CACNA1C* variants can disrupt excitatory-inhibitory balance and network organization, potentially contributing to neurodevelopmental phenotypes and susceptibility to disorders such as epilepsy and ASD.

Acute LTCC blockade with diltiazem largely reproduced known effects on network activity, shortening burst intervals as previously reported in iPSC-derived neurons^20^. We also observed reduced basal firing, not previously reported in this model. However, LTCC blockade has been shown to reduce activity in a seizure model, highlighting the role of LTCCs in neuronal firing^33^. In contrast, *CACNA1C* LoF neurons exhibited both overlapping and distinct phenotypes: while burst intervals were similarly reduced, LoF lines showed increased active electrodes and a higher proportion of spikes within bursts. This suggests chronic loss of Cav1.2 triggers compensatory or developmental changes in addition to direct effects on neuronal firing. The patient line, carrying a novel A1521P variant, showed an opposite phenotype, with reduced excitability, fewer active electrodes, and prolonged burst intervals. indicating the variant has a gain-of-function effect on Cav1.2 channels.

Patients with variants in *CACNA1C* show diverse neurodevelopmental symptoms, including developmental delay, intellectual disability and ASD^7,8^. The effect of channel function can vary across variants, and our cellular models suggest that both gain and loss of function can perturb development, contributing to these phenotypes. In addition to these symptoms, around 40-50% of patients develop seizures or epilepsy ^7,8^. Our findings suggest CACNA1C loss of function may increase seizure susceptibility, whereas the A1521P variant reduced excitability, consistent with the patient’s lack of seizures. In patients, seizures appear to be more common in non-truncating variants of *CACNA1C*, though their effects on channel function vary^8^. This suggests both loss and gain of function may contribute to seizures and nuances in the effect of each variant plays a role in the variable phenotype seen across patients with *CACNA1C* variants. Though our patient has no history of seizures, they do have an ASD diagnosis, and the neuronal phenotypes align with features previously reported in iPSC-based ASD models showing reduced spontaneous excitatory postsynaptic current (sEPSC) frequency, spike rate, synchronised burst frequency, and dynamic complexity of cultures^34–36^.

GABAergic signalling emerged as a central mechanism underlying these network phenotypes. LoF lines showed reduced GABAergic signalling and diazepam resistance, while the GoF line displayed increased inhibition, as indicated by restored activity after GABA blockade. These results align with prior studies showing that LTCC blockade reduces surface expression of GABAA receptors, decreases the number of parvalbumin interneurons and decreases protein expression of GAD65, GAD67 and VGAT ^37–40^. Similarly, heterozygous *CACNA1C* knockout rats show decreased GAD67+ and Parvalbumin+ neurons ^41^. In contrast, LTCC activation leads to increased surface expression of GABA_A_ receptors^42^. TS models also show increases in GABA_A_ receptor expression, GAD67 and VGAT puncta^9,43,44^. Importantly, dysregulation of inhibitory signalling has been implicated in both ASD and epilepsy. In epilepsy, downregulation of GABAergic signalling leads to disinhibition and hyperexcitability, which can then lead to seizures^45^. Both up and down regulation of GABAergic signalling has been associated with ASD in iPSC-derived models and our finding reinforce the idea that either increased or decreased GABAergic tone can destabilise developing networks ^35,46^.

We also observed early developmental alterations that likely contribute to later network phenotypes. *CACNA1C*-related disorder (CRD), ASD and epilepsy all have strong developmental components and LTCCs play an important role in regulation of gene transcription during development. Specifically, Cav1.2 regulates gene transcription via the CREB pathway and LoF and GoF lines showed opposing effects on CREB phosphorylation. These findings are consistent with studies showing reduced CREB signalling in Cav1.2 LoF models and increased signalling in G406R TS models^14,16,47^. These changes provide a potential mechanism for the changes to GABAergic signalling as CREB phosphorylation has been shown to increase GAD67 levels via LTCCs ^39^. Cav1.2 channels also activates CamKII, which is upstream of CREB, leading to increased membrane insertion of GABA_A_ receptors^42^. CREB phosphorylation via LTCCs is important for the generation of neurons from NPCs and CREB is upregulated in NPCs and young neurons ^48^. This may explain the reduced neuron number and increased glial gene expression in LoF lines. Indeed, *CACNA1C*-deficient mouse NPCs generate fewer neurons and more astrocytes and pharmacological manipulation of LTCCs in rodent NPCs alters neuronal-glial balance^49,50^.

Neuron production may also be disrupted by changes to neuronal rosette structure, a critical process in early brain development. Rosettes mimic the radial organization of the neural tube in vitro and are essential for neural patterning and progenitor cell organization. In our model, reduced rosette size in *CACNA1C* LoF lines may reflect impaired cell–cell adhesion. This is consistent with prior evidence that N-cadherin is sensitive to L-type calcium channel activity, and that loss of Cav1.2 in other systems reduces RhoA/ROCK signalling, a key pathway regulating cadherin-mediated adhesion and rosette integrity during neuronal differentiation^51–53^. A reduction in RhoA signalling in CACNA1C LoF lines could compromise adherens junction stability, leading to smaller or less organized rosettes, while the inverse may occur in GoF lines. Consistent with this, RNA-seq data indicated broader downregulation of genes involved in cell junction organization, which could affect cortical architecture at later stages. This includes WNT5A, a gene critical for neuronal migration and cortical layering, and thus may contribute to altered cortical layer specification previously observed in Timothy Syndrome models^9,10,54^.

This study highlights the critical role of Cav1.2 in neurodevelopment, demonstrating its impact on inhibitory–excitatory balance and neuronal network activity. Across models, *CACNA1C* disruption altered CREB signalling, NPC maintenance, and rosette organization, linking early developmental changes to later shifts in excitatory–inhibitory balance and network dynamics. Importantly, the CACNA1C LoF and GoF lines showed contrasting phenotypes: loss of Cav1.2 increased network excitability, while the A1521P variant led to reduced excitability and prolonged burst intervals. Although this work is based on a single patient-derived iPSC line, and we cannot draw conclusions for the wider CRD patient population, this model allows us to investigate the consequences of both loss and gain of function on human neurobiology. Additionally, the contrasting effects between LoF and GoF variants underscore how different classes of *CACNA1C* variants may disrupt neuronal development and function through distinct mechanisms. Given the wide spectrum of *CACNA1C* variants and associated clinical phenotypes, future studies comparing iPSC-derived neurons from multiple patients with diverse *CACNA1C* variants will be essential to define shared pathways and clarify how specific channel alterations translate into the wide spectrum of neurodevelopmental phenotypes observed in patients. Moreover, this work establishes a set of functional readouts—including network activity measures, ERK/CREB pathway activation, and neurodevelopmental patterning phenotypes—that will provide a valuable framework for defining the cellular consequences of *CACNA1C* variants in future studies.

## Acknowledgements and Disclosures

This work was funded by the Medical Research Council (MRC) through the GW4 BioMed Doctoral Training Partnership and by a research fellowship funded by The Waterloo Foundation. Patient-derived samples used to generate induced pluripotent stem cells were obtained as part of the IMAGINE-ID study of individuals with intellectual difficulties of presumed genetic origin, with informed consent and approval from South-East Wales REC.

The authors report no biomedical financial interests or potential conflicts of interest.

