## Supplementary Material for "Gain and loss of function changes in *CACNA1C* affect neuronal networks through divergent pathways"

### Supplemental Methods

##### Human iPSC culture and neuronal differentiation

The cell lines used in this study were human induced pluripotent stem cell lines (iPSCs). The control lines were an IBJ4 human iPSC line derived from the BJ fibroblast cell line (ATCC; CRL-2522), HPSI1013i-wuye_2 (HipSci) and an inducible Cas9 line, which was derived from the IBJ4 line. Cells were cultured on Matrigel (Corning) in Essential 8™ medium (ThermoFisher) at 37°C, 5% $CO_2$ and passaged with Gentle Cell Dissociation Reagent or Accutase (StemCell Technologies). Differentiation into cortical glutamatergic neurons followed a modified protocol by Chambers et al. using dual SMAD inhibition (SB431542, LDN193189) in N2B27 media (2/3 DMEM/F12; 1/3; neurobasal; B27-RA; N2; 1xPSG; 0.1 mM β-mecaptoethanol)[27]. Once NPCs were generated cells were cultured in N2B27 with B27 + retinoic acid on PDL (Sigma)/laminin (Roch) at 200000 cells/cm2. At day 50, neurons were plated onto multi-electrode arrays (MEAs) and maintained in BrainPyhs (Stem Cell Technologies) supplemented with B27+RA, PSG; 10ng/ml BDNF and 200µM ascorbic acid.

##### Generation of *CACNA1C* LoF iPSCs

A CRISPR-Cas9 inducible IBJ4 line (pAAVS1-PDi-CRISPRn; Addgene #73500) was used to target *CACNA1C*. Cas9 expression was induced with doxycycline (2 µg/ml) and cells were transfected with three synthetic gRNAs using Lipofectamine CRISPRMAX. Successful editing was confirmed by PCR amplification using the flanking primers 5‘-GGTGAGGCAAGGAGACTAGAGC-3‘ and 5‘-GATTCATCCAGTCTAGGGCGGG-3’. PCR products were then sequenced using minION long read sequencing (ONT) to identify deletion sizes. Off-target effects were assessed using CRISPOR and RGEN Cas-OFFinder; 12 predicted sites showed no mutations.

##### Generation of patient CACNA1C iPSCs

A patient with a *CACNA1C* p.Ala1521Pro variant was recruited via the IMAGINE-ID study of individuals with intellectual difficulties of presumed genetic origin (ethics approval: South-East Wales REC). A peripheral whole-blood sample was obtained by JFGU. Full details of the IMAGINE-ID protocol have previously been reported[28]. iPSCs were generated from peripheral blood mononuclear cells using the Sendai-based CytoTune-iPS 2.0 kit (A16517, ThermoFisher scientific).

##### Western blot analysis

Cells were lysed in RIPA buffer (R0278, Sigma) and protease inhibitor cocktail (MSSAFE, Sigma) for 30 mins at 4°C. 20 µg of protein per sample was separated on 4-12% Bis-Tris Plus Gels (Life Technologies) and transferred to nitrocellulose. Membranes were blocked in 5% milk in TBST, probed overnight with primary antibodies. Antibodies used were against Cav1.2 (1:200, Millipore, AB5156), ERK1/2 (1:1000, CST, 4695), p-ERK1/2 (1:1000, CST, 9106), CREB (1:1000, CST, 9104), p-CREB (1:1000, CST, 9198) and GAPDH (1:1000, Abcam, ab9485). After washing in TBST blots were incubated with IRDye®- conjugated secondary antibodies (1:10,000, Li-COR). Detection was via Li-COR Odyssey infrared imaging system (Biosciences, Biotechnology). All data normalization was against GAPDH.

##### Immunostaining Analysis

Cells were fixed in 3.7% PFA, permeabilized (0.3% Triton X-100), blocked (5% donkey serum, 1% BSA), and incubated with primary antibodies overnight at 4°C. Cells were washed with PBS before incubation with secondary antibodies and then counterstained with DAPI (1:2000, Molecular Probes) and mounted in ProLong Gold antifade reagent (Invitrogen). Samples were imaged on a Leica DMI6000b fluorescent microscope. An average of 10 images were taken per coverslip. Primary antibodies were as follows: Pax6 (1:1000, AB_528427, DSHB), Ki67 (1:1000, AB15580, Abcam), N-Cadherin (1:1000, 33-3900, Invitrogen), NeuN (1:1000, ab279297, Abcam). Secondary antibodies were as follows: Alexa Fluor 488-conjugated donkey anti-rabbit (1 : 500, A32790, Invitrogen), Alexa Fluor 555-conjugated donkey anti-mouse (1:500, A31570, Invitrogen), Alexa Fluor 647-conjugated donkey anti-chicken (1 : 500, 78952, Invitrogen). Analysis was performed using CellProfiler.

##### Bioinformatic Analysis

RNA sequencing was performed on 3 samples from Control and 3 samples from *CACNA1C* LoF day 30 neurons. Samples were sent to Active motif for library preparation and sequencing. Sequence reads were mapped to the genome using the STAR algorithm. Counts were read into R 4.3.2 (https://www.R-project.org/) and filtered to retain genes with counts greater than 5. One *CACNA1C* LoF sample was deemed an outlier by RUVseq and excluded form analysis. Differential gene expression (DGE) analysis was performed using DESeq2 (R package, v1.30.0). P values were adjusted for multiple comparisons using Benjamini-Hochberg correction address false discovery rate (FDR). A gene was considered differentially expressed if it has an adjusted P-value of less than 0.01 and a Log2 fold change of ±0.5 or greater.

Enrichment analysis was performed to determine Gene Ontology – Biological Processes (GOBP), using the fgsea multilevel enrichment test and genes were ranked by minimum significant difference signed msd (fgsea). Enriched terms with a padj value less than 0.01 were considered significant. Enriched GOBP terms were clustered based on semantic similarity using GOSemSim (R package, v2.24.0), and clusters were manually summarized. Tissue enrichment analysis was conducted using the TissueEnrich (R package, v1.20.0). Disease-gene network analysis was performed using the DisGeNET database (V7.0), calculating overlap significance for DEGs and disease genesets by hypergeometric probability, with adjusted $P<0.001$ considered statistically significant.

##### Gene Expression Analysis

RNA was extracted from cells using the RNeasy mini kit (Qiagen, Germany). For each sample 500ng of RNA was reverse transcribed using the High-Capacity RT cDNA Kit (ThermoFisher Scientific). qRT-PCR analysis of mRNA used the qPCRBIO SyGreen Blue Mix (PCR Biosystems, UK). All qRT-PCR reactions were performed in triplicate on a QuantStudio 7 Flex RT-PCR machine (ThermoFisher Scientific) and relative expression was calculated using the Pfaffl method with data normalised to GAPDH and B-Actin (see Table S1 for primer sequences).

#### MEA and pharmacological manipulation

Day 50 neurons were plated as high density drop cultures (15,000 cells/µl) containing 10 µg/ml laminin onto PEI-treated 24-well CytoView MEAs (M384-tMEA-24 W, 4X4 electrode grid). Cells were cultured in 1:1 fresh to normal human astrocytes (NHA) conditioned BrainPhys media with half-media changes twice a week. Neurons were treated with DAPT (10 µM) for 1 week to remove NPCs and allow synchronised networks to form. Activity was recorded using Axion Maestro Pro. Channels were sampled with a gain of 1000 × and a sampling rate of 12.5 kHz/channel. During the recording, the temperature was maintained at 37 °C. Quality control excluded wells with coverage of less than 80% of electrodes. Pharmacological agents (bicuculline, diltiazem, diazepam) were applied in BrainPhys for 10 min (1 h for diazepam) before recording. After recordings, medium was removed from the MEAs and cultures were washed 3 times with PBS before fresh medium was added.

Spike detection was carried out using AxiS software (Adaptive Threshold Crossing Method, 6 x Standard Deviations). Offline analysis was achieved with custom scripts written in R. Spike timestamps were analysed to provide statistics on the general excitability of cultures. Network activity was analysed using array–wide spike detection rate (ASDR) with a bin width of 200 ms. Synchronised bursts were detected from binned data by a 3-step process: The start of a synchronised burst was detected by a spike count of at least 40% of the maximum ASDR. If the subsequent bin(s) also contained a spike count above the threshold it was included in the synchronised burst. The end of a synchronised burst was determined by a period of 400ms or more without a spike count about the threshold.

#### Statistical Analysis

Data were analysed in GraphPad Prism 8. Normality was assessed via Anderson-Darling test; ROUT (Q=1%) was used to exclude outliers. One-way or two-way ANOVA was applied as appropriate. Data are presented as mean and standard error of the mean (SEM). Statistical significance was considered as P-values <0.05, and significances represented as ∗P < 0.05, ∗∗P < 0.01 and ∗∗∗P < 0.001.

### Supplemental Tables

| Gene | Forward | Reverse |
|---|---|---|
| GAPDH | AGGCTGGGGCTCATTTG | CAGTTGGTGGTGCAGGAG |
| B-Actin | TCACCACCACGGCCGAGC | TCTCCTTCTGCATCCTGTCG |
| GABRA1 | GGATTGGGAGAGCGTGTAACC | TGAAACGGGTCCGAAACTG |
| GABRB2 | TCTCTCTGTATACGATGGACCC | GCTTCTGGGGTCTCCAAGTC |
| GAD65 | GGCTTTTGGTCTTTCGGGTC | GCACAGTTTGTTTCCGATGCC |
| GAD67 | GCCAGACAAGCAGTATGATGT | CCAGTTCCAGGCATTTGTTGAT |
| MAP2 | CTGCTTTACAGGGTAGCACAA | TTGAGTATGGCAAACGGTCTG |
| vGlut1 | CGACGACAGCCTTTTGTGGT | GCCGTAGACGTAGAAAACAGAG |
| vGlut2 | GGGAGACAATCGAGCTGACG | CAGCGGATACCGAAGGAGATG |
| CACNA1C (Exon 2) | GGTGAGGCAAGGAGACTAGAGC | GATTCATCCAGTCTAGGGCGGG |
| CACNA1C (Exons 1-4) | CATTTCTTCCTCTTCGTGGCTGC | TAGTAGGTTCCAGCCGTTGC |

**Table S1: Primers for qRT-PCR analysis.** This table lists the forward and reverse primer sequences used for qRT-PCR analysis for each gene examined.

| Measure | Condition | | | | | | ANOVA | Comparison to Control |
|---|---|---|---|---|---|---|---|---|
| | Control | | LoF | | GoF | | | |
| | Mean | SEM | Mean | SEM | Mean | SEM | | |
| Well Spike Rate (Hz) | 18.05 | 2.12 | 28.26 | 4.24 | **4.775** | 0.9075 | W = 28.39, p < 0.0001 | KO p = 0.0747, **Patient p < 0.0001** |
| Median Electrode Spike Rate (Hz) | 5.94 | 0.71 | 4.755 | 0.6623 | **2.985** | 0.4778 | F = 5.895, p = 0.0039 | KO p = 0.3981, **Patient p = 0.0019** |
| Mean Spike Amplitude (mV) | 0.022 | 0.0010 | 0.025 | 0.0017 | **0.017** | 0.00076 | W = 12.95, p < 0.0001 | KO p = 0.4613, **Patient p = 0.0002** |
| Number Active Electrodes | 10.98 | 0.53 | **13.04** | 0.56 | **7.50** | 0.83 | F = 14.25, p < 0.0001 | **KO p = 0.0317, Patient p = 0.0004** |
| Mean SB interval (s) | 51.22 | 0.78 | **29.21** | 0.44 | **76.77** | 2.84 | KW = 31.88, p < 0.0001 | **KO p = 0.0066, Patient p = 0.0007** |
| SB interval Co-efficient of variation | 0.69 | 0.041 | 1.30 | 0.223 | 0.84 | 0.096 | KW = 1.878, p = 0.3910 | NA |
| Number Synchronised bursts | 42.55 | 1.92 | **69.13** | 3.91 | **32.50** | 3.86 | W = 24.79, p < 0.0001 | **KO p < 0.0001, Patient p = 0.0486** |
| Mean Spikes per SB | 124.5 | 18.50 | 129.10 | 18.89 | **42.56** | 6.89 | W = 15.76, p < 0.0001 | KO p = 0.9816, **Patient p = 0.0002** |
| Percent of spikes in SBs (%) | 40.91 | 2.19 | **53.50** | 2.76 | 48.99 | 3.33 | F = 5.878, p = 0.0038 | **KO p = 0.0034**, Patient p = 0.0612 |

**Table S2: Mean ± SEM and statistical analysis of multielectrode array (MEA) measures.** This table summarizes measures obtained from control, LoF and GoF lines. For each parameter, values are presented as mean ± SEM. Statistical tests were selected based on data distribution: one-way ANOVA was used for normally distributed data, and Kruskal–Wallis tests were used for non-normal data. Post-hoc comparisons were performed using Dunnett’s or Dunn’s test, respectively. Exact p-values are reported, with significance defined as p < 0.05. Data from at least 3 independent differentiations across 2-3 cell lines per group.

### Supplemental Figures

#### Figure S1

A

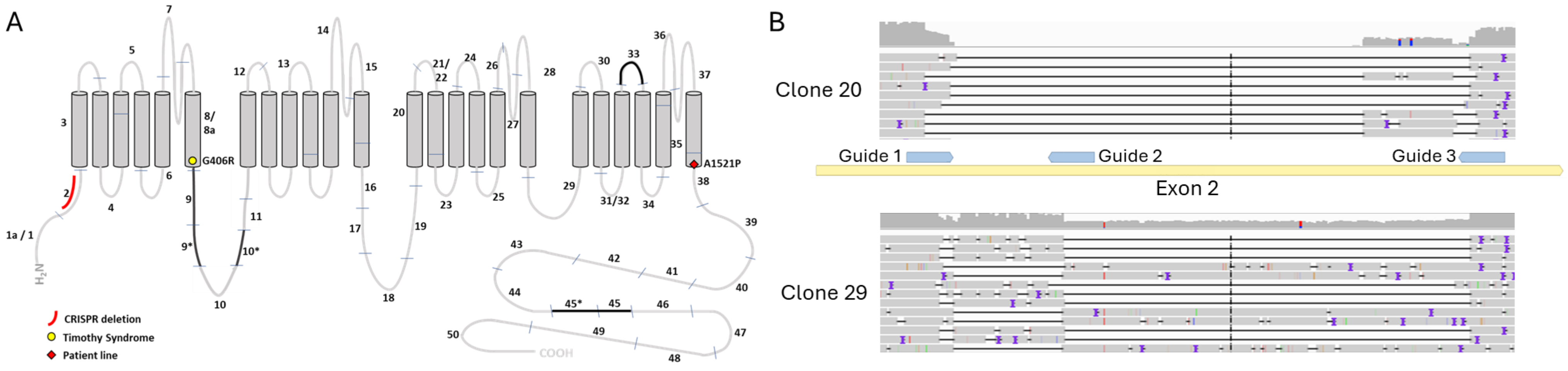


C

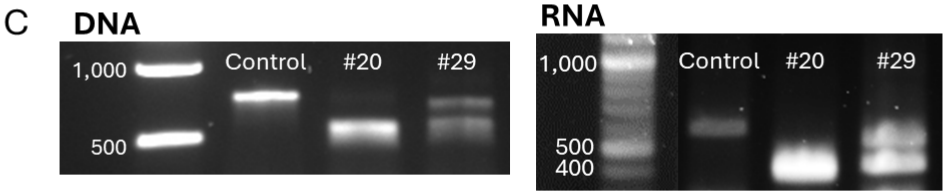


D

| Clone | INDELs | |
|---|---|---|
| 20 | 224 bp deletion | 183 + 8bp deletion |
| 29 | 6 bp + 176 bp deletion | 47 bp deletion |

E

**Wild-type**

GSNYGSPRPAHANM ... PIRRACISIVEWKPFEIIILLT...

**Deletions**

GSNYGSPRPAHANM ... PHPEGLHQHCRMETI *

F

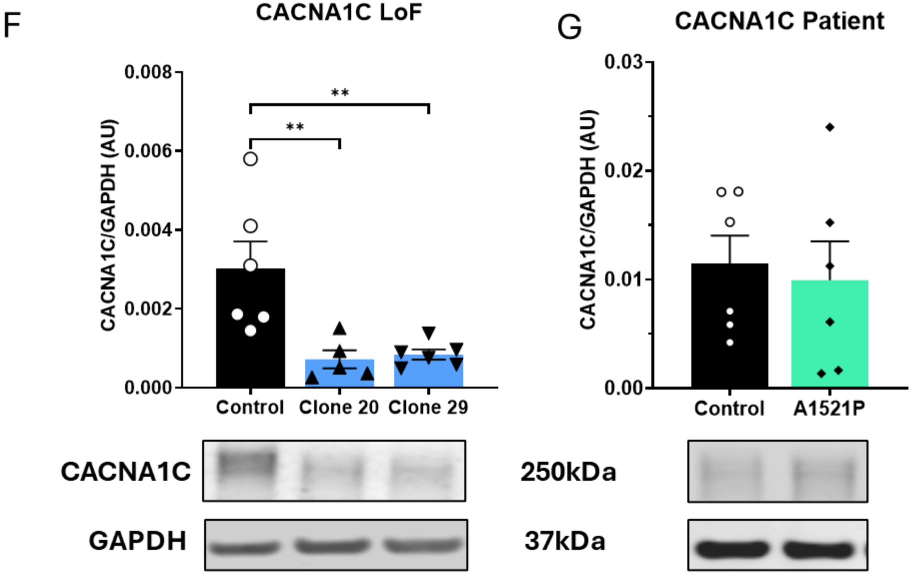

**Figure S1: Generation of CACNA1C LoF lines and validation of protein expression in CACNA1C LoF and Patient A1521P lines A:** Diagram of CACNA1C structure showing location of typical Timothy Syndrome mutation, deletion in LoF lines and point mutation in patient line. **B:** Sequencing results of the two clones that contained indels. Each row is an individual read from the sequencing and the black lines denote deletions when compared to the reference genome. The position of the guides on exon 2 is shown and is the same scale as the sequencing data. **C:** PCR product of DNA and RNA transcript for target region in iCas9 iPSCs (control) and the two clones that contained indels ([illegible]20 and [illegible]29). **D:** Summary of deletion sizes present in clones. **E:** Deletions in exon 2 of CACNA1C LoF lines leads to introduction of stop codon in exon 3 of CACNA1C. **F:** Western Blot of CACNA1C with GAPDH as housekeeping in iCas9 and CACNA1C LoF lines at day 50 of neuronal differentiation **G:** Western Blot of CACNA1C with GAPDH as housekeeping in control and patient A1521P lines at day 50 of neuronal differentiation. Plots show mean ± SEM, individual wells are overlayed as points. *$p < 0.05$, **$p < 0.01$, *** $p < 0.001$, **** $p < 0.0001$ following One-Way ANOVA for Control vs LoF and T-test for Control vs Patient. Data from 2 independent differentiations.

Figure S2:

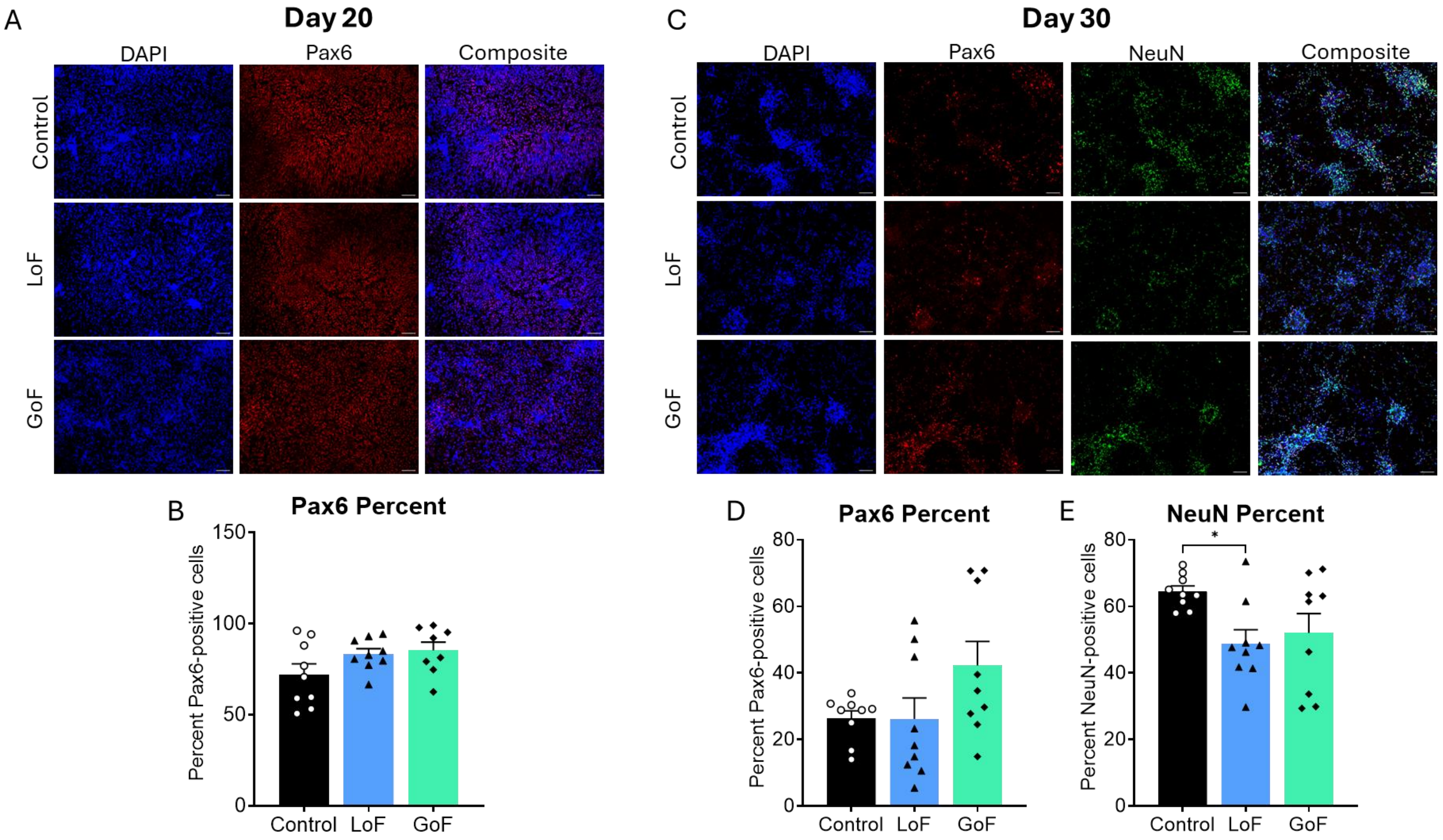

A
Day 20
DAPI
Pax6
Composite
Control
LoF
GoF
C
Day 30
DAPI
Pax6
NeuN
Composite
Control
LoF
GoF
B
Pax6 Percent
Percent Pax6-positive cells
150
100
50
0
Control
LoF
GoF
D
Pax6 Percent
Percent Pax6-positive cells
80
60
40
20
0
Control
LoF
GoF
E
NeuN Percent
Percent NeuN-positive cells
*
80
60
40
20
0
Control
LoF
GoF

**Figure S2: Neuronal differentiation markers in CACNA1C LoF and GoF lines. A:** Examples images of Pax6 staining at day 20 of neuronal differentiation in control, CACNA1C LoF and CACNA1C GoF lines. **B:** Quantification of the percentage of DAPI+ nuclei that colocalized with Pax6 at day 20. **C:** Examples images of Pax6 and NeuN staining at day 30 of neuronal differentiation in control, CACNA1C LoF and CACNA1C GoF lines. **D:** Quantification of the percentage of DAPI+ nuclei that colocalized with Pax6 at day 30. **E:** Quantification of the percentage of DAPI+ nuclei that colocalized with NeuN at day 30. All plots show mean ± SEM, individual replicates are overlayed as points. *$p < 0.05$, **$p < 0.01$, *** $p < 0.001$ following One Way ANOVA. Data from at least 3 independent differentiations across 2 cell lines per group.
